# Phylogenetic Analysis Of SARS-CoV-2 In The First Months Since Its Emergence

**DOI:** 10.1101/2020.07.21.212860

**Authors:** Matías J. Pereson, Laura Mojsiejczuk, Alfredo P. Martínez, Diego M. Flichman, Gabriel H. Garcia, Federico A. Di Lello

**Author notes:** Corresponding author: Dr. Federico Alejandro Di Lello, Facultad de Farmacia y Bioquímica, Universidad de Buenos Aires, Instituto de Investigaciones en Bacteriología y Virología Molecular (IBaViM). Junín 956, 4º piso, (1113), Ciudad Autónoma de Buenos Aires, Argentina.

## Abstract

During the first months of SARS-CoV-2 evolution in a new host, contrasting hypotheses have been proposed about the way the virus has evolved and diversified worldwide. The aim of this study was to perform a comprehensive evolutionary analysis to describe the human outbreak and the evolutionary rate of different genomic regions of SARS-CoV-2.

The molecular evolution in nine genomic regions of SARS-CoV-2 was analyzed using three different approaches: phylogenetic signal assessment, emergence of amino acid substitutions, and Bayesian evolutionary rate estimation in eight successive fortnights since the virus emergence.

All observed phylogenetic signals were very low and trees topologies were in agreement with those signals. However, after four months of evolution, it was possible to identify regions revealing an incipient viral lineages formation despite the low phylogenetic signal, since fortnight 3. Finally, the SARS-CoV-2 evolutionary rate for regions nsp3 and S, the ones presenting greater variability, was estimated to values of 1.37 × 10^−3^ and 2.19 × 10^−3^ substitution/site/year, respectively.

In conclusion, results obtained in this work about the variable diversity of crucial viral regions and the determination of the evolutionary rate are consequently decisive to understand essential feature of viral emergence. In turn, findings may allow characterizing for the first time, the evolutionary rate of S protein that is crucial for vaccines development.

## Introduction

Coronaviruses belong to *Coronaviridae* family and have a single strand of positive-sense RNA genome of 26 to 32 kb in length ^[1]^. They have been identified in different avian hosts as well as in various mammals including bats, mice, dogs, etc. ^[2,3]^. Periodically, new mammalian coronaviruses are identified. In late December 2019, Chinese health authorities identified groups of patients with pneumonia of unknown cause in Wuhan, Hubei Province, China ^[4]^. The pathogen, a new coronavirus called SARS-CoV-2 ^[5]^, was identified by local hospitals using a surveillance mechanism for “pneumonia of unknown etiology” ^[4,6,7]^. The pandemic has spread rapidly and, to date, more than 22 million confirmed cases and nearly 750,000 deaths have been reported in just over a six months period ^[8]^. This rapid viral spread raises interesting questions about the way its evolution is driven during the pandemic. From the SARS-CoV-2 genome, 16 non-structural proteins (nsp1-16), 4 structural proteins [spike (S), envelope (E), membrane (M) and nucleoprotein (N)], and other proteins essential to complete the replication cycle are translated ^[9,10]^. The large amount of information currently available allows knowing, as never before, the real-time evolution history of a virus since its interspecies jump ^[11]^. Most studies published to date have characterized the viral genome and evolution by analyzing complete genomes sequences ^[12,13,14,15]^. Despite this, until now, the viral genomic region providing the most accurate information to characterize SARS-CoV-2, could not be established. This lack of information prevent from investigating its molecular evolution and monitoring biological features affecting the development of antiviral and vaccines. Therefore, the aim of this study was to perform a comprehensive viral evolutionary analysis in order to describe the human outbreak and the molecular evolution rate of different genomic regions of SARS-CoV-2.

## Materials and Methods

### Datasets

In order to generate a dataset representing different geographic regions and time evolution of the SARS-CoV-2 pandemic from December 2019 to April 2020, data of all the complete genome sequences available at GISAID (https://www.gisaid.org/) on April 18, 2020 were collected. Data inclusion criteria were: a.-complete genomes, b.-high coverage level, and c.-human hosts only (no other animals, cell culture, or environmental samples). Complete genomes were aligned using MAFFT against the Wuhan-Hu-1 reference genome (NC_045512.2, EPI_ISL_402125). The resulting multiple sequence alignment (dataset 1) was split in nine datasets corresponding to nine coding regions: a.-four structural proteins [envelope (E), nucleocapsid (N), spike (S), Orf3a], b.-four nonstructural proteins (nsp1, nsp3, Orf6, and nsp14), and c.-an unknown function protein (Orf8).

More than six thousand SARS-CoV-2 publicly available nucleotide sequences were downloaded. After data selection according to the inclusion criteria, 1616 SARS-CoV-2 complete genomes were included in dataset 1. Sequences of this dataset 1 came from 55 countries belonging to the five continents as follow: Africa: 39 sequences, Americas: 383 sequences, Asia: 387 sequences, Europe: 686 sequences and Oceania: 121 sequences. After elimination of sequences with indeterminate or ambiguous positions, the number of analyzed sequences for each region were: nsp1, 1608; nsp3, 1511; nsp14, 1550; S, 1488; Orf3a, 1600; E, 1615; Orf6, 1616; Orf8, 1612; and N, 1610. Finally, nucleotide sequences were grouped by fortnight (FN) according to their collection date. Table 1 summarizes the number of sequences per fortnight since the beginning of the pandemic up to FN 8. On the other hand, Dataset 2 was created using only variable sequences of each region analyzed in Dataset 1. Thus, Dataset 1 was used for the analysis of amino acid substitutions and Dataset 2 was used for the phylogenetic signal analysis and the Bayesian coalescent trees construction.

**Table 1.** Number of SARS-CoV-2 sequences by fortnight (Temporal structure)

| Fortnight | Date | Median of analyzed sequences<br>(Q1-Q3) |
| --- | --- | --- |
| FN1 | 12/24/2019 to 12/31/2019 | 15 |
| FN2 | 01/01/2020 to 01/15/2020 | 19 |
| FN3 | 01/16/2020 to 01/31/2020 | 145 (136-145.5) |
| FN4 | 02/01/2020 to 02/15/2020 | 119 (113-120) |
| FN5 | 02/16/2020 to 03/02/2020 | 258 (247-259) |
| FN6 | 03/03/2020 to 03/17/2020 | 403 (390-406) |
| FN7 | 03/18/2020 to 04/01/2020 | 447 (416-450) |
| FN8 | 04/02/2020 to 04/17/2020 | 199 (197-201) |
| **TOTAL** |  | **1488 to 1616** |

FN: Fortnight; Q1=quartile 1, Q3=quartile 3. The total number of sequences is variable depending on the analyzed region (nsp1, 1608; nsp3, 1511; nsp14, 1550; S, 1488; Orf3a, 1600; E, 1615; Orf6, 1616; Orf8, 1612; and N, 1610)

### Phylogenetic signal

To determine the phylogenetic signal of each of the nine generated alignments, Likelihood Mapping analyzes were carried out ^[16]^, using the Tree Puzzle v5.3 program ^[17]^ and the Quartet puzzling algorithm. This algorithm allowed analyzing the tree topologies that can be completely solved from all possible quartets of the n alignment sequences using maximum likelihood. An alignment with defined tree values greater than 70-80% presents strong support from the statistical point of view ^[17]^. Identical sequences were also removed with ElimDupes (Available at https://www.hiv.lanl.gov/content/sequence/elimdupesv2/elimdupes.html) as they increase computation time and provide no additional information about data phylogeny. The best-fit evolutionary model to each dataset was selected based on the Bayesian Information Criterion obtained with the JModelTest v2.1.10 software ^[18]^.

### Analysis of amino acid substitutions

Entropy-One (Available at https://www.hiv.lanl.gov/content/sequence/ENTROPY/entropy_one.html) was used to determining in dataset 1 the frequency of amino acids at each position for the nine genomic regions analyzed and evaluating their permanence in the eight investigated fortnights.

### Bayesian coalescence and phylogenetic analysis

To study the relationship between SARS-CoV-2 sequences, nine regions of the virus genome were investigated by Bayesian analyses. Phylogenetic trees were constructed using Bayesian inference with MrBayes v3.2.7a ^[19]^. Each gene was analyzed independently with the same dataset used for the phylogenetic signal analysis so that non-identical sequences were included in the analysis. Analyses were run for five million generations and sampled every 5000 generations. Convergence of parameters [effective sample size (ESS) ≥ 200, with a 10% burn-in] was verified with Tracer v1.7.1 ^[20]^. Phylogenetic trees were visualized with FigTree v1.4.4.

### Evolutionary rate

The estimation of the nucleotide evolutionary rate was made with the Beast v1.10.4 program package ^[21]^. Analyses were run at the CIPRES Science Gateway server ^[22]^. Three hundred and twelve sequences without indeterminations corresponding to the nsp3 (5835nt) and S (3822nt) genes were randomly selected from dataset 1. The sequences represent all the fortnights and most of the geographical locations sampled until April 17. Temporal calibration was performed by date of sampling. The appropriate evolutionary model was selected as described above for phylogenetic signal analysis. The TIM model of nucleotide substitution was used for nsp3 and, the HKY model of nucleotide substitution for S. The analysis was carried out under a relaxed (uncorrelated lognormal) molecular clock model suggest by Duchene & col. ^[23]^ and with an exponential demographic, proper for early viral samples from an outbreak ^[24]^. Independent runs were performed for each dataset and a Markov chain Monte Carlo (MCMC) with a length of 1.3×10^9^ steps, sampling every 1.3×10^6^ steps, was set. The convergence of the “meanRate” parameters [effective sample size (ESS) ≥ 200, burn-in 10%] was verified with Tracer v1.7.1 ^[20]^. Additionally, in order to verify the obtained results, 15 independent replicates of the analysis were performed with the time calibration information (date of sampling) randomized as described by Rieux & Khatchikian, 2017 ^[25]^. Finally, the obtained parameters for real data and the randomized replicates were compared.

## Results

### Phylogenetic signal

Using bioinformatics tools, a phylogenetic signal study was carried out in order to identify the most informative SARS-CoV-2 genomic regions. The likelihood mapping analysis showed that most genes has very poor phylogenetic signal with high values in central region which represents the area of unresolved quartets (Figure 1). Accordingly, genes could be separated into three groups. A group with little or no phylogenetic signal (E, Orf6, Orf8, nsp1, and nsp14), a second group with low phylogenetic signal (Orf3a and N), and a last group with relatively more phylogenetic signal (S and nsp3) but still low to be considered a robust one (unresolved quartets >40%).

**Figure 1.**
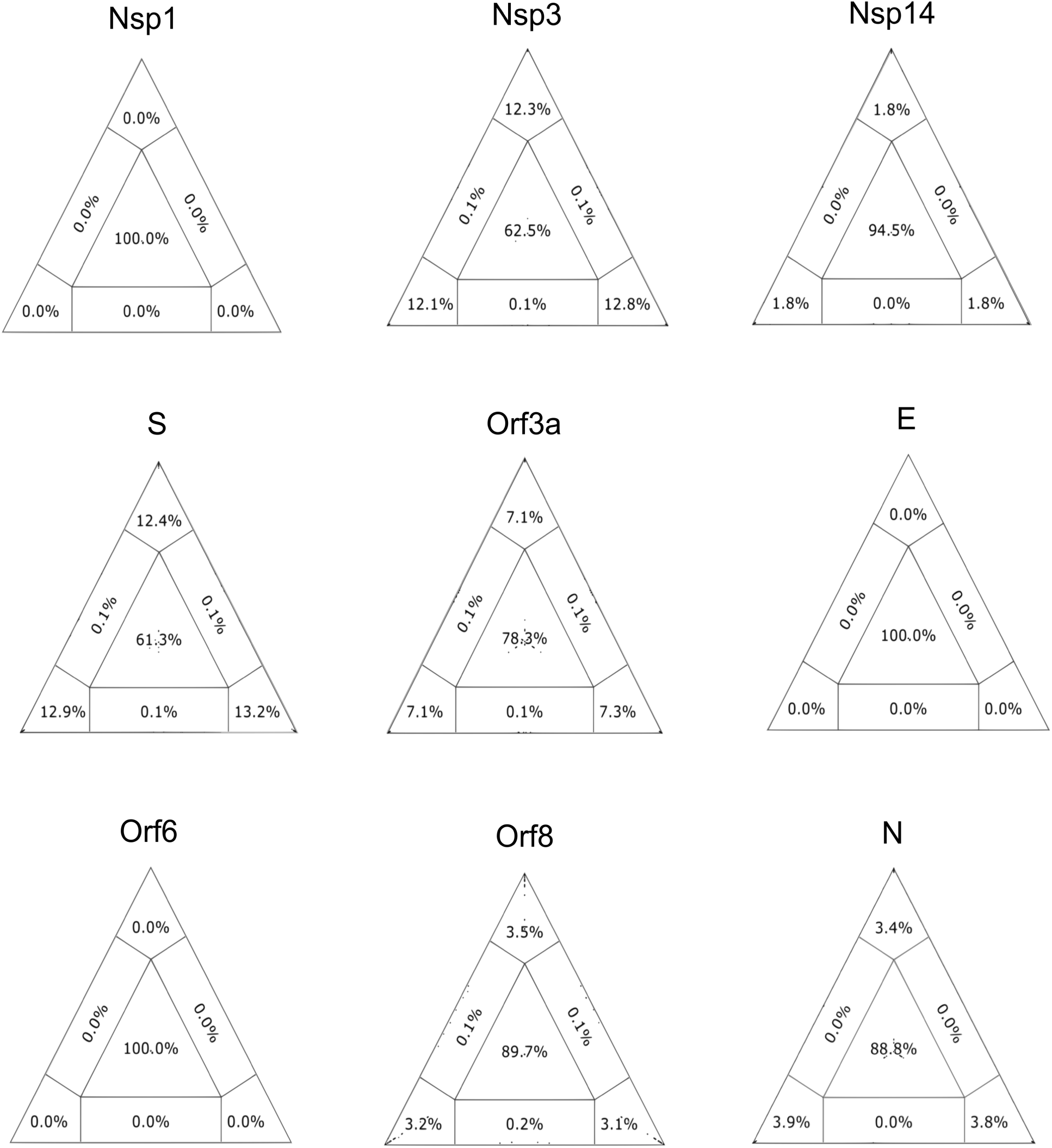
Phylogenetic signal for SARS-CoV-2 datasets. Presence of phylogenetic signal was evaluated by likelihood mapping, unresolved quartets (center) and partly resolved quartets (edges) for genomes available on April 17 for the nine analyzed regions: nsp1 (29 sequences), nsp3 (225 sequences), nsp14 (65 sequences), S (183 sequences), Orf3a (74 sequences), E (11 sequences), Orf6 (12 sequences), Orf8 (23 sequences), and N (113 sequences). Presence of strong phylogenetic signal (<40% unresolved quartets) was not reached for any region.

### Analysis of amino acid substitutions

The analysis of amino acids substitutions by fortnights was useful to study the viral evolutionary dynamics in the context of the beginning of the pandemic. By analyzing different time periods amino acid sequences, changes were observed in 5 out of 9 genomic regions and only in 14 out of the 4975 (0.28%) evaluated residues. In most of the regions, except nsp1, nsp14, E, and Orf6, 2 to 6 amino acids were selected since FN3 and remain unchanged until the end of the follow up period (Table 2). Particularly, in Orf8 region, early selection of two amino acid substitutions (V62L and L84S) was observed from FN2. On the other hand, in the S region, the D614G substitution started with less than 2% in FN3 and FN4 and reached 88% in the last fortnight. In a similar way, the Q57H (Orf3a) substitution went from 6% to 34% while L84S (Orf8) start to be selected in FN2 and reached 6% by FN8. The R203K and G204R substitutions of the N region was selected in FN4 and increased their population proportion with values greater than 20% towards the end of the follow up period. Moreover, selection of a great number of sporadic substitutions remaining in the population for a short period (1-3 fortnights) was observed in the nine analyzed regions. Indeed, 333 (6.83%) of the analyzed positions presented at least one substitution throughout the eight fortnights. Table 3 summarizes the number of variable positions, number of mutations, and number of sequences with mutations by region.

**Table 2.** Amino acids selected by region and fortnight. The number indicates the amino acids location in its protein.

| Region | Amino acid substitution | Amino acid percentage by FN |  |  |  |  |  |  |  |
| --- | --- | --- | --- | --- | --- | --- | --- | --- | --- |
|  |  | FN1 | FN2 | FN3 | FN4 | FN5 | FN6 | FN7 | FN8 |
| nsp3 | A58T | 0 | 0 | 0 | 1.0 | 6.0 | 3.0 | 3.0 | 2.5 |
|  | P135L | 0 | 0 | 0.8 | 0 | 0 | 1.5 | 0.5 | 2.5 |
| S | D614G | 0 | 0 | 1.5 | 1.8 | 37.0 | 64.0 | 75.0 | 88.0 |
| Orf3a | Q75H | 0 | 0 | 0 | 0 | 6.0 | 22.0 | 23.0 | 34.0 |
|  | G196V | 0 | 0 | 0 | 0 | 0.8 | 4.0 | 0.9 | 0.5 |
|  | G251V | 0 | 0 | 8.0 | 24.0 | 8.0 | 9.0 | 10.0 | 3.0 |
| Orf8 | V62L | 0 | 5.0 | 1.0 | 3.3 | 0.0 | 1.5 | 1.3 | 3.0 |
|  | L84S | 0 | 42.0 | 37.0 | 21.0 | 21.0 | 18.0 | 7.0 | 6.0 |
| N | P13L | 0 | 0 | 0 | 0 | 1.0 | 1.0 | 2.5 | 0.5 |
|  | S197L | 0 | 0 | 0 | 0 | 1.1 | 5.0 | 0.9 | 0.5 |
|  | S202N | 0 | 0 | 3.5 | 4.2 | 0 | 0.5 | 2.2 | 2.5 |
|  | R203K | 0 | 0 | 0 | 0 | 17.0 | 19.0 | 24.0 | 23.0 |
|  | G204R | 0 | 0 | 0 | 0 | 17.0 | 19.0 | 24.0 | 23.0 |
|  | I292T | 0 | 0 | 0 | 0 | 2.0 | 0.2 | 0.2 | 0.5 |

Only regions where amino acid change was selected and remained until the last analyzed fortnight are shown. FN: Fortnight; aa: amino acid

**Table 3.** Number of variable positions, number of mutations, and number of sequences with mutation by region

| **Region** | **Nº of variable aa**<br>**positions (%)** | **Nº of aa**<br>**substitutions** | **Nº of sequences with**<br>**aa substitutions (%)** |
| --- | --- | --- | --- |
| nsp1 (180aa) | 3 (1.7) | 37 | 37 (2.4) |
| nsp3 (1945aa) | 158 (8.1) | 322 | 294 (19.3) |
| nsp14 (527aa) | 6 (1.4) | 83 | 83 (5.5) |
| S (1273aa) | 76 (5.9) | 1013 | 904 (59.4) |
| Orf3a (275aa) | 11 (4) | 491 | 468 (30.7) |
| E (75aa) | 5 (6.7) | 6 | 6 (0.4) |
| Orf6 (60aa) | 7 (11.6) | 9 | 9 (0.6) |
| Orf8 (121aa) | 14 (11.6) | 312 | 288 (18.9) |
| N (419aa) | 53 (12.6) | 760 | 470 (30.9) |
| Total (4875aa) | 333 (6.8) | 3033 | - |

aa: amino acid

### Bayesian coalescence analysis

In this study, trees were performed by Bayesian analysis instead of by distance, likelihood, or parsimony methods. Consistently with the phylogenetic signal analysis, trees for nsp1, E, and Orf6 showed a star-like topology. Nevertheless, different proportions of clades formation could be observed in trees of Orf8, nsp14, Orf3a, N, S, and nsp3 regions (Figure 2). Finally, from mentioned regions, nsp3 and S showed a better clade constitution. This analysis allowed to differentiate regions presenting a diversification process (nsp3, nsp14, Orf3a, S, Orf8, and N) from those that even after four months showed an incipient one (nsp1, E, and Orf6). Furthermore, this nucleotide analysis is complemented by the previous study of amino acid variations in each region. However, it is important to note that due to the low phylogenetic signal observed for each region, results can only be considered as preliminary.

**Figure 2.**
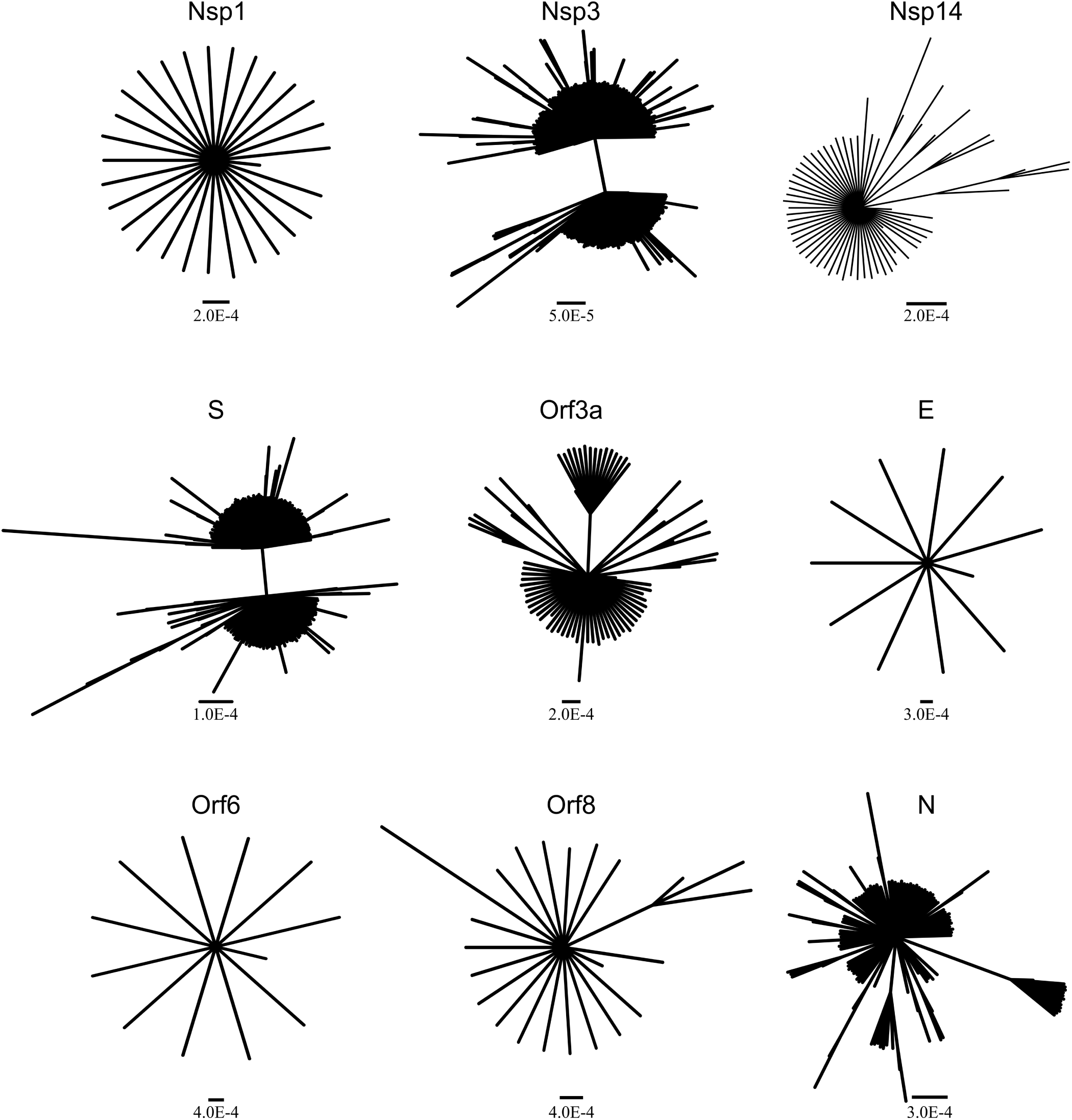
Bayesian trees of 29 sequences of nsp1 (540nt), 225 sequences of nsp3 (5835nt), 65 sequences of nsp14 (1581nt), 183 sequences of S (3822nt), 74 sequences of Orf3a (828nt), 11 sequences of E (228nt), 12 sequences of Orf6 (186nt), 23 sequences of Orf8 (366nt), and113 sequences of N (1260nt). Scale bar represents substitutions per site.

### Evolutionary rate

Nsp3 and S sequences were selected to perform the evolutionary rate analysis since both regions provided the best phylogenetic information among studied regions. The observed evolutionary rate for nsp3 protein of SARS-CoV-2 was estimated to be 1.37 x10^−3^ (ESS 782) nucleotide substitutions per site per year (s/s/y) (95% HPD interval 9.16 x10^−4^ to 1.91 x10^−3^). On the other hand, the corresponding figures for S were estimated to be 2.19 x10^−3^ (ESS 383) nucleotide s/s/y (95% HPD interval 3.19 x10^−3^ to 1.29 x10^−3^). In both genomic regions, date-randomization analyses showed no overlapping between the 95% HPD substitution-rate intervals obtained from real data and from date-randomized datasets. This fact suggests that the original dataset has enough temporal signal to perform analyses with temporal calibration based on tip-dates (Figure 3).

**Figure 3.**
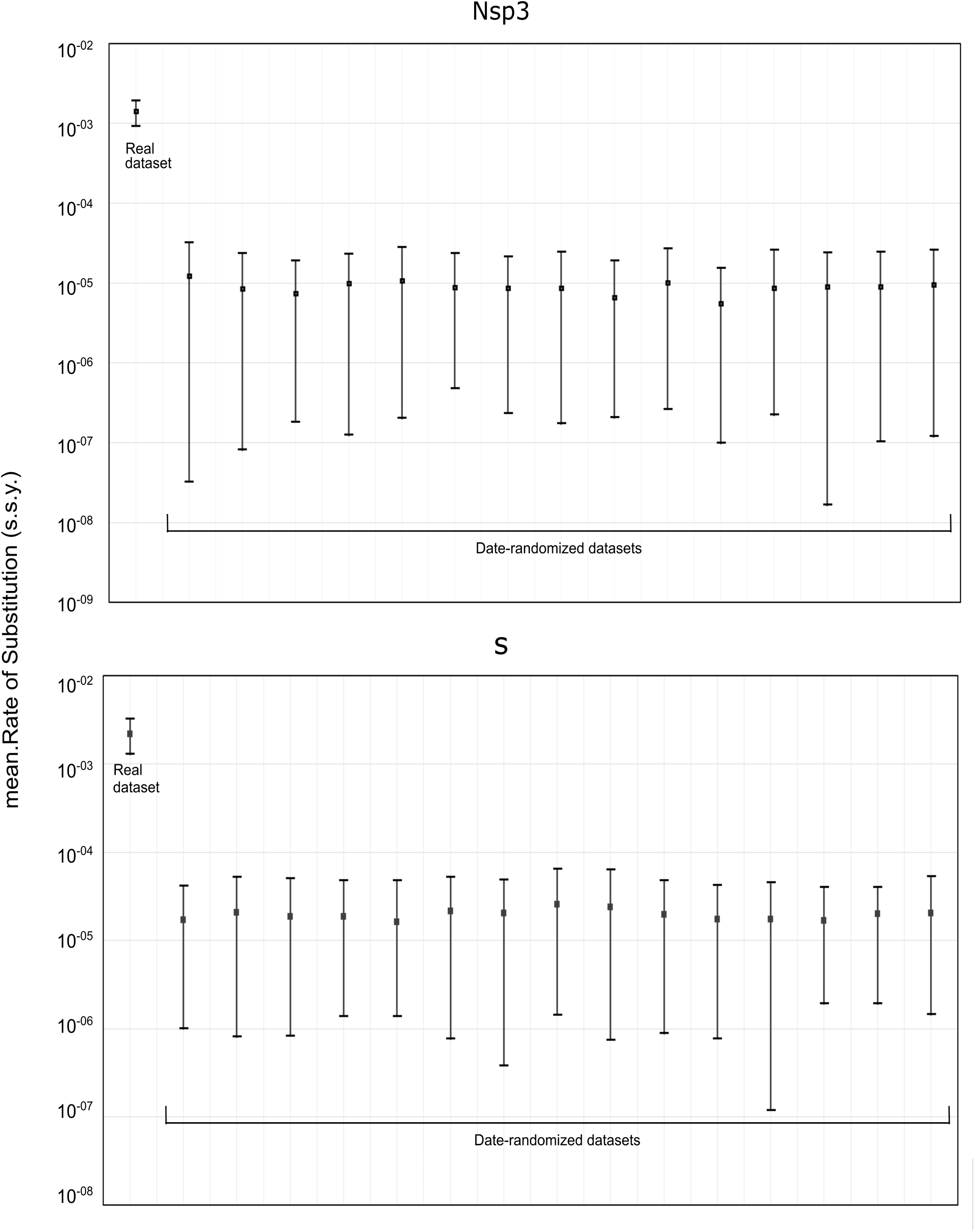
Comparison of the evolutionary rates estimated using BEAST for the original dataset and the date-randomized datasets (312 sequences). This analysis was performed for regions nsp3 (5835nt) and S (3822nt). s.s.y = substitutions/site/year.

## Discussion

The phylogenetic characterization of an emerging virus is crucial to understand the way the virus and the pandemic will evolve. Thereby, a detailed study of the SARS CoV-2 genome allows, on the one hand, to contribute to the knowledge of viral diversity in order to detect the most suitable regions to be used as antivirals or vaccines targets. On the other hand, the large amount of information that is continuously generated, is allowing studying the SARS CoV-2 genome and describing a new viral real time evolution like never before.

In the present study, the molecular evolution and viral lineages of SARS-CoV-2 in nine genomic regions, during eight successive fortnights, was analyzed using three different approaches: phylogenetic signal assessment, emergence of amino acid substitutions, and Bayesian evolutionary rate estimation. In this context, the observed phylogenetic signals of nine coding regions were very low and the obtained trees were consistent with this finding, showing star-like topologies in some viral regions (nsp1, E, and Orf6). However, after a four months evolution period, it was possible to identify regions (nsp3, S, Orf3a, Orf8, and N) revealing an incipient formation of viral lineages, despite the phylogenetic signal, both at the nucleotide and amino acid levels from FN3. Based on these findings, the SARS-CoV-2 evolutionary rate was estimated, for the first time, for the two regions showing higher variability (S and nsp3).

As regards the phylogenetic signal, several simulation studies has proven that for a set of sequences to be considered robust, the central and lateral areas representing the unresolved quartets, must not be greater than 40% ^[16]^. In this regard, none of the nine analyzed regions met this requirement. Three regions (E, nsp1, and Orf6) presented values of 100% unresolved quartets. Most regions (nsp14, Orf3a, Orf8, and N) reached values higher than 85%. Only in regions nsp3 and S the number of unresolved quartets dropped to ∼ 60%. Thus, despite being a virus with an RNA genome, the short time elapsed since its emergence, and possibly genetic restrictions have led to a constrained evolution of SARS-CoV-2 in these months. For this reason, it is expected that trees generated from SARS-CoV-2 partial sequences in the first months of the pandemic are unreliable for defining clades. Therefore, they should be analyzed with great caution.

Since Bayesian analysis allows to infer phylogenetic patterns from tree distributions, it represents a more reliable tool to compare different evolutionary behaviors. Bayesian analysis helps to obtain a tree topology that is closer to reality in the current conditions of SARS-CoV-2 pandemic ^[26]^. The phylogenetic analysis for nsp1, E, and Orf6 regions confirmed the star-like topologies in accordance to a lower diversification of these regions using the sequences available up to FN8 (Figure 2). Trees generated from nsp14 and Orf8 are at an intermediate point, where the formation of small clusters can be observed. In fact, a mutation at position 28,144 (Orf8: L84S) has been proposed as a possible marker for viral classification ^[27,28]^. Finally, trees obtained from regions Orf3a, N, nsp3, and S showed the best clade formation. Indeed, in the most variable regions nsp3 and S, it can be clearly seen that sequences are separated into two large groups. Despite the aforementioned for the nsp3 and S regions, even clusters with very high support values should be taken with precaution and longer periods should be considered to obtain more accurate phylogeny data. However, even when data are not the most accurate to study the spread or clade formation ^[29, 30]^, they provide a good representation of the way the virus is evolving.

The analysis of amino acids frequencies allowed identifying different degree of region conservation throughout the viral genome as a consequence of positive and negative pressures. In particular, nsp3, S, Orf8, and N showed some substitutions in high frequencies. This would indicate, as other authors previously report, the frequent circulation of polymorphisms due to significant positive pressure ^[13,27,31]^. Additionally, since S and N are among candidates to be used in the formulation of vaccines and antibody treatment, it will be important to monitor these substitutions in different geographic regions in order to improve treatment and vaccination efficacy ^[32,33,34]^. In particular, the appearance of the D614G variant in the third week and its rapid increase until reaching a prevalence of 88% in the eighth week could reflect an improvement in viral fitness, as several studies reported ^[35]^.

Contrarily, in regions nsp1, nsp14, E, and Orf6 no substitutions were selected and lasted during the first 4 months of the pandemic. This would suggest that these are regions with constraints to change due to the great negative selection pressure, as it has been recently reported ^[13]^.

In the present study, the evolutionary rate for SARS-CoV-2 genes was estimated by analyzing a large number of sequences, which were carefully curated and had a good temporal and spatial structure. Additionally, the most phylogenetically informative regions of the genome (nsp3 and S) were used for analysis, reinforcing the results confidence. Previous studies on SARS-CoV-2 have reported similar data ranging from 1.79 × 10^−3^ to 6.58 × 10^−3^ s/s/y for the complete genome ^[6,36]^. However, in both articles, small datasets of complete genomes were used (N=32 and 54, respectively). As studies were performed early in the outbreak and due to datasets temporal structure, analysis could have led to less precise estimates of the evolutionary rate ^[23]^. On the other hand, another study from van Dorp et al. (2020), analyzing 7,666 sequences has obtained different results with a remarkably low evolutionary rate (6 × 10^−4^ nucleotide/genome/year) ^[15]^. However, it is important to consider that van Dorp et al. (2020) estimate the evolutionary rate using the complete genome, including several highly conserved genomic regions, while in our work, the estimation was performed with the most variable regions of the genome. Additionally, tests randomizing the dates of nsp3 and S datasets were carried out; they showed that these partial genomic regions have enough temporal signal. In this context, our results (1.37 × 10^−3^ s/s/y for NSp3 and 2.19 × 10^−3^ s/s/y for S) are in close agreement with those published for SARS-CoV genome, which have been estimated between 0.80 to 3.01 × 10^−3^ s/s/y ^[37-39]^(The Chinese SARS Molecular Epidemiology Consortium, 2004, Vega et al. 2004, Zhao et al. 2004). Moreover, our values are in the same order magnitude as other RNA viruses ^[40]^. Even though we should be cautious with these results interpretation, the date-randomization analysis indicated a robust temporal signal.

In addition, the importance of separately studying the evolutionary rate in S region arises from the fact that it represents the main target for antiviral agents and vaccines since it includes the SARS-CoV-2 binding receptor domain (RBD), a crucial structure for the virus to enter host cells and binding site for neutralizing antibodies ^[41]^.

Despite limitations of the evolutionary study of an emerging virus, where the selection pressures are still low and therefore its variability is also low, this work has a great strength: it lies on the extremely careful selection of a big sequence dataset to be analyze. First, it was considered selected sequences to have a good temporal signal and spatial (geographic) structure. Secondly, much attention was paid to the elimination of sequences with low coverage and indeterminacies that could generate a noise for the phylogenetic analysis of a virus that is beginning to evolve in a new host.

The appearance of a new virus means an adaptation challenge. The SARS-CoV-2 overcome the spill stage and shows a significantly higher spread than SARS-CoV and MERS-CoV, thus becoming itself the most important pandemic of the century. In this context, the results obtained in this work about the variable diversity of nine crucial viral regions and the determination of the evolutionary rate, are consequently decisive to understanding essential feature of viral emergence. Nevertheless, monitoring SARS-CoV-2 population will be required to determine the evolutionary course of new mutations as well as to understand the way they affect viral fitness in human hosts.

## Competing interest

On behalf of all authors, the corresponding author states that there is no conflict of interest.

## Funding

None

## Declaration of Author Contributions

MJP: Data curation, acquisition of data, analysis and interpretation of data, drafting the article, final approval of the version to be submitted.

LM: Data curation, acquisition of data, analysis and interpretation of data, revising the article critically for important intellectual content, final approval of the version to be submitted.

APM: Data curation, Validation, revising the article critically for important intellectual content, final approval of the version to be submitted.

DMF: Data curation, Validation, drafting the article, final approval of the version to be submitted.

GG: Data curation, acquisition of data, analysis and interpretation of data, drafting the article, final approval of the version to be submitted.

FAD: Conception and design of the study, acquisition of data, analysis and interpretation of data, drafting the article, final approval of the version to be submitted.

## Acknowledgements

MJP, LM, DMF, and FAD are members of the National Research Council (CONICET). We would like to thank to the researchers who generated and shared the sequencing data from GISAID (https://www.gisaid.org/) and Mrs. Silvina Heisecke from CEMIC-CONICET for providing language assistance.

## Notes

### Competing Interest Statement

The authors have declared no competing interest.

## REFERENCES

[1] Su S, Wong G, Shi W, et al. Epidemiology, genetic recombination, and pathogenesis of coronaviruses. Trends in Microbiology 2016; 24, 490–502. https://doi.org/10.1016/j.tim.2016.03.003

[2] Cavanagh D. Coronavirus avian infectious bronchitis virus. Veterinary Research 2007; 38, 281–297. https://doi.org/10.1051/vetres:2006055

[3] Ismail MM, Tang AY & Saif YM. Pathogenicity of turkey coronavirus in turkeys and chickens. Avian Diseases 2003; 47, 515–522. https://doi.org/10.1637/5917

[4] Zhu N, Zhang D, Wang W, et al. A Novel Coronavirus from Patients with Pneumonia in China, 2019. The New England Journal of Medicine 2020; 382, 727–733. https://doi.org/10.1056/NEJMoa2001017

[5] Coronaviridae Study Group of the International Committee on Taxonomy of Viruses. The species Severe acute respiratory syndrome-related coronavirus: classifying 2019-nCoV and naming it SARS-CoV-2. Nature Microbiology 2020; 5, 536–544. https://doi.org/10.1038/s41564-020-0695-z

[6] Li X, Wang W, Zhao X, et al. Transmission dynamics and evolutionary history of 2019-nCoV. Journal of Medical Virology 2020a; 92, 501–511. https://doi.org/10.1002/jmv.25701

[7] Li Q, Guan X, Wu P, et al. Early Transmission Dynamics in Wuhan, China, of Novel Coronavirus-Infected Pneumonia. The New England Journal of Medicine 2020b; 382, 1199–1207. https://doi.org/10.1056/NEJMoa2001316

[8] World Healt Organization, 2020. Coronavirus disease (COVID-19) Situation Report –118. Retrieved from: https://www.who.int/docs/default-source/coronaviruse/situation-reports/20200517-covid-19-sitrep-118.pdf?sfvrsn=21c0dafe_6 (15 August 2020, date last accessed).

[9] Cui J, Li F & Shi ZL. Origin and evolution of pathogenic coronaviruses. Nature Reviews Microbiology 2019; 17, 181–192. https://doi.org/10.1038/s41579-018-0118-9

[10] Luk HKH, Li X, Fung J, et al. Molecular epidemiology, evolution and phylogeny of SARS coronavirus. Infection Genetics and Evolution 2019; 71, 21–30. https://doi.org/10.1016/j.meegid.2019.03.001

[11] Zhou P, Yang XL, Wang XG, et al. A pneumonia outbreak associated with a new coronavirus of probable bat origin. Nature 2020; 579, 270–273. https://doi.org/10.1038/s41586-020-2012-7

[12] Benvenuto D, Giovanetti M, Salemi M, et al. The global spread of 2019-nCoV: a molecular evolutionary analysis. Pathogens and Global Health 2020; 114, 64–67. https://doi.org/10.1080/20477724.2020.1725339

[13] Cagliani R, Forni D, Clerici M, et al. Computational inference of selection underlying the evolution of the novel coronavirus, SARS-CoV-2. Journal of Virology 2020; https://doi.org/10.1128/JVI.00411-20

[14] Phan T. Genetic diversity and evolution of SARS-CoV-2. Infection Genetics and Evolution 2020; 81, 104260. https://doi.org/10.1016/j.meegid.2020.104260

[15] van Dorp L, Acman M, Richard D, et al. Emergence of genomic diversity and recurrent mutations in SARS-CoV-2. Infection Genetics and Evolution 2020; 5, 104351. https://doi.org/10.1016/j.meegid.2020.104351

[16] Strimmer K & von Haeseler A. Likelihood-mapping: A simple method to visualize phylogenetic content of a sequence alignment. Proceedings of the National Academy of Sciences of the USA 1997; 94, 6815–6819. https://doi.org/10.1073/pnas.94.13.6815

[17] Schmidt HA, Strimmer K, Vingron M, et al. TREE-PUZZLE: Maximum likelihood phylogenetic analysis using quartets and parallel computing. Bioinformatics 2002; 18, 502–504. https://doi.org/10.1093/bioinformatics/18.3.502

[18] Darriba D, Taboada GL, Doallo R, et al. jModelTest 2: more models, new heuristics and parallelcomputing. Nature Methods 2012; 9, 772. https://doi.org/10.1038/nmeth.2109

[19] Ronquist F, Teslenko M, van der Mark P, et al. MrBayes 3.2: efficient Bayesian phylogenetic inference and model choice across a large model space. Systematic Biology 2012; 61, 539–542. https://doi.org/10.1093/sysbio/sys029

[20] Rambaut A, Drummond AJ, Xie D, et al. Posterior summarization in Bayesian phylogenetics using Tracer 1.7. Systematic Biology 2018; 67, 901–904. https://doi.org/10.1093/sysbio/syy032

[21] Suchard MA, Lemey P, Baele G, et al. Bayesian phylogenetic and phylodynamic data integration using BEAST 1.10. Virus Evolution 2018; 4, vey016. https://doi.org/10.1093/ve/vey016

[22] Miller MA, Pfeiffer X, & Schwartz T. Creating the CIPRES Science Gateway for inference of large phylogenetic trees. Gateway Computing Environments Workshop 2010; 1–8. https://doi.org/10.1109/GCE.2010.5676129

[23] Duchene S, Featherstone L, Haritopoulou-Sinanidou M, et al. Temporal signal and the phylodynamic threshold of SARS-CoV-2. bioRxiv 2020; [Preprint]. https://doi.org/10.1101/2020.05.04.077735

[24] Grassly NC & Fraser C. Mathematical models of infectious disease transmission. Nat Rev Microbiol 2008; 6, 477–487. https://doi.org/10.1038/nrmicro1845

[25] Rieux A & Khatchikian CE. tipdatingbeast: an r package to assist the implementation of phylogenetic tip-dating tests using beast. Molecular Ecology Resources 2017; 17, 608–613. https://doi.org/10.1111/1755-0998.12603

[26] Drummond AJ, Ho SY, Phillips MJ, et al. Relaxed phylogenetics and dating with confidence. PLoS Biology 2006; 4, e88. https://doi.org/10.1371/journal.pbio.0040088

[27] Tang X, Wu C, Li X, et al. On the origin and continuing evolution of SARS-CoV-2. National Science Review 2020; 0, 1–12. https://doi.org/10.1093/nsr/nwaa036

[28] Yin C. Genotyping coronavirus SARS-CoV-2: methods and implications. Genomics 2020; 30318–30319. https://doi.org/10.1016/j.ygeno.2020.04.016

[29] Mavian C, Marini S, Prosperi M, et al. A snapshot of SARS-CoV-2 genome availability up to 30th March, 2020 and its implications. JMIR Public Health Surveill 2020; 6, e19170.https://doi.org/10.2196/19170

[30] Sánchez-Pacheco SJ, Kong S, Pulido-Santacruz P, et al. Median-joining network analysis of SARS-CoV-2 genomes is neither phylogenetic nor evolutionary. Proceedings of the National Academy of Sciences of the USA 2020; 117, 9241–9243. https://doi.org/10.1073/pnas.2007062117

[31] Issa E, Merhi G, Panossian B, et al. S.SARS-CoV-2 and ORF3a: Nonsynonymous Mutations, Functional Domains, and Viral Pathogenesis. mSystems 2020 [Preprint]. https://doi.org/10.1128/mSystems.00266-20

[32] Ahmed SF, Quadeer AA & McKay MR. Preliminary Identification of Potential Vaccine Targets for the COVID-19 Coronavirus (SARS-CoV-2) Based on SARS-CoV Immunological Studies. Viruses 2020; 12, 254. https://doi.org/10.3390/v12030254

[33] Callaway E. The race for coronavirus vaccines: a graphical guide. Nature 2020; 580, 576–577. https://doi.org/10.1038/d41586-020-01221-y

[34] Koyama T, Weeraratne D, Snowdon JL, et al. Emergence of Drift Variants That May Affect COVID-19 Vaccine Development and Antibody Treatment. Pathogens 2020; 9, 324. https://doi.org/10.20944/preprints202004.0024.v1

[35] Li Q, Wu J, Nie J, et al. The Impact of Mutations in SARS-CoV-2 Spike on Viral Infectivity and Antigenicity. Cell 2020; S0092-8674(20)30877-1. Advance online publication. https://doi.org/10.1016/j.cell.2020.07.012

[36] Giovanetti M, Benvenuto D, Angeletti S, et al. The first two cases of 2019-nCoV in Italy: Where they come from? Journal of Medical Virology 2020; 92, 518–521. https://doi.org/10.1002/jmv.25699

[37] The Chinese SARS Molecular Epidemiology Consortium. Molecular Evolution of the SARS Coronavirus During the Course of the SARS Epidemic in China. Science 2004; 303, 1666–1669. https://doi.org/10.1126/science.1092002

[38] Vega VB, Ruan Y, Liu J, et al. Mutational dynamics of the SARS coronavirus in cell culture and human populations isolated in 2003. BMC Infectious Diseases 2004, 4, 32. https://doi.org/10.1186/1471-2334-4-32

[39] Zhao Z, Li H, Wu X, et al. Moderate mutation rate in the SARS coronavirus genome and its implications. BMC Evolutionary Biology 2004; 4, 21. https://doi.org/10.1186/1471-2148-4-21

[40] Sanjuán R. From molecular genetics to phylodynamics: evolutionary relevance of mutation rates across viruses. PLoS Pathogens 2012; 8, e1002685. https://doi.org/10.1371/journal.ppat.1002685

[41] Ju B, Zhang Q, Ge J, et al. Human neutralizing antibodies elicited by SARS-CoV-2 infection. Nature 2020; 115–119. https://doi.org/10.1038/s41586-020-2380-z

